# The Effect of Hybridization on Dosage Compensation in Member Species of the *Anopheles gambiae* Species Complex

**DOI:** 10.1101/327577

**Authors:** Kevin C. Deitz, Willem Takken, Michel A. Slotman

**Affiliations:** Department of Entomology, Texas A&M University, College Station, TX 77843, USA; Current address: Department of Ecology and Evolutionary Biology and The Lewis-Sigler Institute for Integrative Genomics, Princeton University, Princeton, NJ 08544, USA; Department of Plant Sciences, Laboratory of Entomology, Wageningen University, Wageningen, The Netherlands

**Keywords:** *Anopheles*, Dosage Compensation, Hybridization, RNAseq

## Abstract

Dosage compensation has evolved in concert with Y-chromosome degeneration in many taxa that exhibit heterogametic sex chromosomes. Dosage compensation overcomes the biological challenge of a "half dose" of X chromosome gene transcripts in the heterogametic sex. The need to equalize gene expression of a hemizygous X with that of autosomes arises from the fact that the X chromosomes retain hundreds of functional genes that are actively transcribed in both sexes and interact with genes expressed on the autosomes. Sex determination and heterogametic sex chromosomes have evolved multiple times in Diptera, and in each case the genetic control of dosage compensation is tightly linked to sex determination. In the *Anopheles gambiae* species complex (Culicidae), maleness is conferred by the Y-chromosome gene *Yob*, which despite its conserved role between species is polymorphic in its copy number between them. Previous work demonstrated that male *An. gambiae s.s.* males exhibit complete dosage compensation in pupal and adult stages. In the present study we have extended this analysis to three sister species in the *An. gambiae* complex: *An. coluzzii*, *An. arabiensis*, and *An. quadriannulatus*. In addition, we analyzed dosage compensation in bi-directional F1 hybrids between these species to determine if hybridization results in the mis-regulation and disruption of dosage compensation. Our results confirm that dosage compensation operates in the *An. gambiae* species complex through the hyper-transcription of the male X chromosome. Additionally, dosage compensation in hybrid males does not differ from parental males, indicating that hybridization does not result in the mis-regulation of dosage compensation.

## Introduction

Sex chromosomes evolved independently from autosomes multiple times within Diptera (Vicoso and Bachtrog, 2015, Bachtrog *et al.*, 2014). While X chromosomes largely retain their original gene content (Vicoso and Charlesworth, 2006), sexually antagonistic mutations that occur on nascent Y-chromosomes suppress recombination with the X (Rice 1996). This leads to Y-chromosome degeneration as male-biased genes move from the X to the Y chromosome or the autosomes or are lost (Parisi *et al.*, 2003), and eventually the formation of heterogametic sex chromosomes (Bachtrog 2013). Old Y-chromosomes harbor few protein coding genes and are repeat-rich, hampering, until recently, their sequencing and characterization (Mahajan and Bachtrog, 2017).

As neo-Y chromosomes degenerate, X-linked genes that are not female-biased in their expression must be equalized or compensated for in the heterogametic sex. The evolution of dosage compensation (DC) has occurred in concert with Y-chromosome degeneration in many taxa that exhibit heterogametic sex determination (Disteche 2012, Graves 2015). DC overcomes the biological challenge of a "half dose" of X chromosome gene transcripts (or Z chromosome transcripts in ZZ/ZW systems) in the heterogametic sex. The need to equalize gene expression of a hemizygous X with that of autosomes arises from the fact that the X chromosomes retain hundreds of functional genes that are actively transcribed in both sexes.

In *Drosophila* and *Anopheles* the regulation of DC is closely tied to the sex-determination pathway despite striking differences in sex determination between these genera. In *Drosophila* females, a double dose of X chromosome-linked signal elements (XSE) initiates female-specific pre-mRNA splicing of *sex lethal* (*Sxl*) transcripts. *Sxl* in turn regulates sex-specific splicing of *transformer* (*tra*), which, along with its non-sex-specific co-factor *transformer2* (*tra2*), promotes female-specific splicing of *doublesex* (*dsx*) and *fruitless* (*fru*) pre-mRNAs. Male and female isoforms of *dsx* and *fru* modulate the expression of genes involved in sexually dimorphic morphologies and physiologies. A single dose of XSE fails to initiate the cascade of female-specific splicing and male transcripts are produced (Penalva and Sanchez, 2003). While male and female isoforms of DSX share the same DNA-binding domain, they differ in their C-terminal domains that confer sex-specific gene regulation (Clough *et al.*, 2014).

*Dsx* and *fru* are highly conserved in insects despite their variety of sex-determination mechanisms (Herpin and Schartl, 2015, Salz and Erickson, 2010, Wilkins 1995). In mosquitoes (Diptera: Culicidae), sex determination is controlled by a M-factor located on the Y chromosome in the subfamily *Anophelinae*, which has heterogametic sex chromosomes (Krzywinska *et al.*, 2016), or on the homomorphic sex-determining chromosome of the subfamily *Culicinae* (Hall *et al.,* 2016). Transcription of the M-factor during early embryo development activates a cascade of sex-specific gene splicing that controls male development through sex-specific splicing of *dsx* and *fru*.

In the *Anopheles gambiae* species complex the M-factor *Yob* is located within the male-determining locus of the Y chromosome*. Yob* expression is male-specific, begins within 2.5 hours of embryo oviposition, and occurs prior to *dsx* splicing by up to six hours (Krzywinska *et al.*, 2016). This indicates that *Yob* is either a direct or indirect upstream regulator of *dsx* splicing. *Yob* is the only protein-coding gene on the Y chromosome that is shared by all members of the *An. gambiae* species complex (Hall *et al.,* 2016). *Yob* is polymorphic in copy number between *An. gambiae* complex species and within *An. gambiae s.s.* (Hall *et al.* 2016)*. An. stephensi*, which diverged from the *An. gambiae* species complex ~30 million years ago, lacks a *Yob* homolog, indicating a recent origin of *Yob* in the *An. gambiae* complex lineage (Hall *et al.* 2016).

Bernardini *et al.* (2017) were able introgress the *An. gambiae* Y chromosome into an *An. arabiensis* genetic background using an F1 × F0 crossing strategy, which allowed them to overcome a severe bottleneck associated with this cross. They found that when isolated, *An. gambiae* and *An. arabiensis* Y-chromosomes are interchangeable; the *An. gambiae* Y chromosome has no effect on *An. arabiensis* gene expression and its introgression does not cause male sterility. Thus, by itself a heterospecific Y-chromosome does not appear to disrupt dosage compensation in this cross. However, we do not know how dosage compensation is affected in F1 hybrids when the genetic background of the autosomes is not homozygous.

DC of the hemizygous X in male *Drosophila* occurs through the binding of the male-specific lethal (MSL) complex to the X. The MSL complex is comprised of four proteins encoded by the male-specific lethal (*msl)* genes 1-3 and *maleless* (*mle)*, all of which cause male inviability when mutated (Penalva and Sanchez, 2003). *Sxl* mutations, or genes impacting its regulation, result in over-expression of genes on the X chromosome. This results in female lethality during embryogenesis (Erickson and Quintero, 2007, Lucchesi and Skripsky, 1981, Gergen 1987, Hilfiker and Amrien, 1995) likely due to the mis-regulation of DC (Biedler and Tu, 2016, Cline 1978).

Like *Drosophila, An. stephensi* and *An. gambiae s.s* exhibit complete DC through the hyper-transcription of the X chromosome (Jing *et al.*, 2015, Rose *et al.,* 2016). Ectopic injections of *Yob* mRNA into *An. gambiae* embryos causes female lethality due to abnormal X chromosome transcription, but has no effect on males. Additionally, knockdown of *Yob* in male *An. gambiae* embryos results in 100% mortality (Krzywinska *et al.*, 2016). Krzywinska *et al* (2016) suggested that these results implicate *Yob* as an upstream regulator of DC in *Anopheles gambiae* due to the interconnected roles of sex-determination genes and DC regulatory pathways in other insects. Orthologs of the five *Drosophila* genes that encode the MSL complex are present in the *An. gambiae* transcriptome. However, Anopheles *msl-1* and *msl-2* are highly diverged from their *Drosophila* orthologs (Rose *et al.*, 2016), which may indicate that they may have different functions in *Anopheles*.

Studies in *Drosophila* initially implicated defects in DC regulation in hybrid male lethality (Chatterjee *et al.*, 2007, Rodriguez *et al.*, 2007, Bachtrog 2008) under a model where MSL proteins and their binding sites evolved on the *D. melanogaster* lineage in a way that a heterospecific mix of MSL complex proteins in F1 *D. melanogaster*-*D. simulans* hybrid males did not effectively regulate DC. Barbash (2010) subsequently refuted this hypothesis through a genetic screen of *mle* and *msl* mutant lines.

The *An. gambiae* species complex harbors the most important vectors of human malaria and has recently emerged as a model system for the study of speciation genetics (Neafsey *et al.*, 2015). However, little is known about dosage compensation regulation in *Anopheles*: sex determination differs drastically between *Drosophila* and *Anopheles,* our knowledge of the link between *Yob* expression and DC regulation is incomplete, and *msl-1* and *msl-2* are highly diverged in sequence, and perhaps function, between *Drosophila* and *Anopheles*.

In the present study, we explored whether DC regulation has diverged between members of *An. gambiae* species complex, and is mis-regulated in their hybrids, through an analysis of genome-wide gene expression in three sister species: *An. coluzzii, An. arabiensis*, and *An. quadriannulatus,* as well as reciprocal F1 hybrids derived from *An. coluzzii* x *An. arabiensis* and *An. coluzzi* x *An. quadriannulatus* crosses.

## Materials and Methods

### Mosquito Rearing

Mosquitoes were derived from the SUA2La (*An. coluzzii*), SANGQUA (aka SANGWE, *An. quadriannulatus*), and DONGOLA (*An. arabiensis*) strains, or from bi-directional crosses between *An. coluzzii* and *An. arabiensis*, as well as *An. coluzzii* and *An. quadriannulatus*. Cross directions are indicated by female x male. Mosquitoes were reared in 12 hr light / 12 hr dark cycles at 25 ° and 70-80% relative humidity with access to 5% sucrose solution. Colonies were blood fed via an artificial membrane with warmed, defibrinated sheep blood (Hemostat Laboratories, Dixon, CA). Adults were provided wetted filter paper during the period of 48-72 hr post-feeding for oviposition. Larvae were fed Tetra brand TetraColor Tropical Crisps (fish food) *ad libitum* until pupation. Mosquitoes were collected 12 hr post-pupation and sexed prior to RNA extraction. Bi-directional crosses were performed by sexing 50-100 male and female pupae from the respective parental colonies prior to mixing them in adult cages. Rearing conditions of F1 larvae and collection of F1 pupae were performed as described above.

### RNA Isolation and Sequencing

Mosquitoes were sexed 12 hr post-pupation. Total RNA extractions were performed on five pupae per replicate using a Qiagen RNeasy Mini kit (Qiagen, Redwood City, CA). RNA samples were quantified using a NanoDrop spectrophotometer and their quality was assessed using an Agilent Bioanalyzer. Equimolar amounts of two to four extraction products (10 or 20 total mosquito pupae) were combined to form a single biological replicate. Pooled RNA samples were again run on the Agilent Bioanalyzer and NanoDrop spectrophotometer prior to library prep and sequencing. Library prep was performed using an Illumina TruSeq RNA library prep kit (Illumina Inc., San Diego, CA). Two biological replicates for each strain, cross, and sex (28 libraries total) were sequenced on seven lanes of Illumina HiSeq 2500 machine using 125 bp single-end chemistry. Raw RNA sequencing reads are available on the NCBI Sequence Read Archive at project accession number PRJNAXXXXXX.

### Pseudo-Genome Construction

The first step of our analysis was to construct an *An. gambiae*-based pseudo-genome for each parental species. By incorporating SNPs from each species into the AgamP4 genome, we were able to use the AgamP4 coordinates and gene set as a common framework to quantify transcript abundance. FastQC package version 0.11.4 (Babraham Bioinformatics) was used to examine read quality. Illumina TruSeq adapters were removed using Trimmomatic version 0.30 (Bolger *et al.*, 2014), while simultaneously soft-clipping the reads from both 5’ and 3’ ends to an average phred quality score of 20, with no single bp in a four bp window below a phred quality score of 10. Only reads ≥ 50 bp were retained for alignment and subsequent analyses.

Trimmed RNAseq data from the three species were initially aligned to their respective, species-specific genomes, which were obtained from VectorBase.org (AcolM1 for *An. coluzzii*, AaraD1 for *An. arabiensis*, and AquaS1 for *An. quadriannulatus*, Giraldo-Calderon *et al.*, 2015). Initial alignments were performed using the splice-aware alignment software STAR version 2.4.2a using default parameters with the exception of the option "--outFilterMismatchNoverLmax 0.1" (Dobin *et al.*, 2013). Next, an index was created for each alignment, which annotated all splice junctions found during the first alignment round. This index informed the second alignment with STAR (as above), which re-aligned the reads around splice junctions. Picard tools version 2.8.1 (http://broadinstitute.github.io/picard/) was used to mark and remove PCR duplicates.

BAM alignments from each of the three species (two biological replicates of males and females) were then merged using Picard MergeBamAlignment, and SAMtools/BCFtools version 1.3 (Li *et al.*, 2009) was used to call variants within each merged alignment using the BCFtools "call" tool. This process was informed by a prior mutation rate of 0.02. Next, GATK was used to filter variant call files for only bi-allelic SNPs. The vcfutils.pl varFilter function, a component of the BCFtools package, was used to filter SNPs using default parameters with the exception of not filtering for strand bias, as this can be expected in RNAseq data.

Filtered SNPs were converted to the genomic coordinates of the *An. gambiae s.s.* PEST strain AgamP4 genome (Holt *et al.*, 2002) using the Picard Tools (github.com/broadinstitute/picard) LiftoverVCF function and a chain file to match base-pair coordinates between genomes. Chain files were created from pair-wise multiple-alignment files between the *An. coluzzii, An. arabiensis*, and *An. quadriannulatus* genomes and the *An. gambiae s.s.* PEST reference originally published by Fontaine *et al.* (2015), which were downloaded from VectorBase.org. Once SNPs were converted to AgamP4 coordinates, they were incorporated into the AgamP4 genome using the GATK FastaAlternateReferenceMaker to create a pseudo-genome for each species using the AgamP4 backbone/coordinates.

Next, we again performed a STAR 2-pass alignment (as above) for each parental library using the species-specific, AgamP4-baesed pseudo-genome as a reference. SNPs were called using the methods described above, and were then incorporated into the previous pseudo-genome reference to further populate the AgamP4-based pseudo-genome of each species with variation from that strain. Subsequently, the AgamP4-based pseudo-genomes were used as a reference to create a cDNA pseudo-genome for each species using the generate_transcripts function of rSeq version 0.2.2 (Jiang *et al.*, 2009, Salzman *et al.*, 2011). Gene transcript coordinates were derived from the AgamP4.4 gene set (VectroBase.org).

### Quantification of Gene Expression

Parental species libraries (*An. coluzzii, An. arabiensis*, and *An. quadriannulatus*) were aligned to their respective cDNA pseudo-genomes using the --very-sensitive mode of Bowtie 2 version 2.2.9 with default parameters (Langmead and Salzberg, 2012). Only uniquely mapped reads were retained to calculate transcript abundances.

RNAseq reads from F1 hybrids were aligned to diploid pseudo-genomes comprised of the AgamP4.4 based cDNA pseudo-genomes of each parental species. rSeq version 0.2.2 (Jiang *et al.*, 2009, Salzman *et al.*, 2011) was used to create cDNA genomes from the AgamP4 based pseudo-genomes of each parental species (as above). The maternal and paternal cDNA genomes for each hybrid were then merged into a diploid genome using the ASE-TIGAR program (Nariai *et al.*, 2016, http://nagasakilab.csml.org/ase-tigar/), which concatenates the cDNA genomes of the maternal and paternal strains into one genome file. RNAseq reads from F1 hybrids were assessed for quality and trimmed using FastQC and Trimmomatic as described above. Reads were mapped to the diploid, bi-parental cDNA pseudo-genomes for the respective cross using the--very-sensitive mode Bowtie 2 version 2.2.9 with default parameters (Langmead and Salzberg, 2012). F1 hybrid reads were allowed to align up to 100 times across the diploid reference genome. ASE-TIGAR was used to identify the most likely parental genome of origin and location of each transcript. Thus, only the highest probability map position of each transcript was quantified. Total transcript abundance (maternal + paternal expression in F1 hybrids) was used for the subsequent analysis of dosage compensation.

### Analysis of Dosage Compensation

Transcript abundances were standardized by converting them to RPKMs (reads per kilo-base of transcript per million reads) in R (www.cran.r-project.org, R Core Team, 2013). Two methods were used to assess dosage compensation in species and F1 hybrid samples. First, the ratio of median expression of X-linked and autosomal genes in each sample was determined.

This was performed by dividing the RPKM value of each X-linked gene by the median RPKM expression level of all autosomal loci. The "boot" package in R (Davison and Hinkley, 1997, Cantly and Ripley, 2015) was used to calculate 95% confidence intervals of the median of this distribution for each sample using 10,000 replicates. X to autosomal (X:A) median gene expression was analyzed at increasing minimum RPKM values between zero and 10. In order to be included in the analysis at a specific RPKM threshold, a gene had to exceed the specified RPKM level in all samples in a species comparison (ex. *An. coluzzii*, *An. arabiensis*, and their hybrids). Because of this, the number of genes included in this analysis differs for *An. coluzzi* between the *An. coluzzi-An. arabiensis* and the *An. coluzzi* - *An. quadriannulatus* species comparisons. An analysis of X:A median gene expression without RPKM cut-off (RPKM ≥ 0.0) would also include transcriptionally inactive genes which would bias the analyses (Kharchenko *et al.*, 2011). Additionally, work in *Drosophila* has shown that genes with low expression have a lower degree of dosage compensation (McAnally and Yampolsky, 2010). Thus, by analyzing X:A expression ratios at increasing minimum RPKM thresholds (RPKM > 0.0, 0.2, 0.5, 1.0, 2.0, 5.0, 10.0) we were able to identify the minimum RPKM level that removes bias during subsequent gene expression analyses.

A comparison of median X:A gene expression ratios between males and females shows if genes on the male hemizygous X chromosome is hyper-expressed to match the level of X chromosome expression in females. In females we expect X:A expression ratios to be at or near one. If the X chromosome is hyper-expressed in males, the X:A expression ratio should be similar to that of females. If dosage compensation is absent, the X:A expression ratio of males should be half that of females. X:A expression ratios in males and females were also compared to the expression ratio of the second and third chromosomes (2:3). We expect the 2:3 expression ratio to be equal in both sexes. Thus, 2:3 median expression serves as an internal standard with which to compare X:A median expression. The 2:3 expression ratios were calculated by correcting the RPKM value of each chromosome 2 gene by the median RPKM of all chromosome 3 genes. The 2:3 expression ratios were also analyzed at increasing minimum RPKM thresholds (RPKM > 0.0, 0.2, 0.5, 1.0, 2.0, 5.0, 10.0).

The 95% confidence intervals of the median of the expression ratio distribution was calculated using the "boot" package in R using 10,000 replicates (Davison and Hinkley, 1997, Cantly and Ripley, 2015). A Kruskal-Wallis test was used to test for significant differences between the median of the X:A expression ratio distribution between each sample (Kruskal and Wallis, 1952). We used an ANOVA to test for significant differences in the mean of the X:A expression ratio distribution within and between each sample in a species comparison. The same tests were performed to analyze significant differences between the distributions of 2:3 expression ratios between samples. Where appropriate, a Tukey’s post-hoc test (Tukey 1949) was performed to test for significant differences in the means of pair-wise comparisons, and a Dunn’s test (Dunn 1964) was used to test for significant differences in the medians of pair-wise comparisons.

Additionally, we calculated male to female (M:F) expression ratios of X-linked and autosomal genes. Genes were only included in this analysis if they exceeded a RPKM of 10.0 across all samples in a comparison (see results). The expectation of this approach is that the M:F ratio of X-linked and the M:F ratio of autosomal genes will not differ if dosage compensation is operating effectively. M:F expression ratios were calculated by first calculating the mean RPKM expression of each gene between the two biological replicates for each sample. Then, the male to female expression ratio was calculated for each gene within a species or F1 hybrid. Medians and 95% confidence intervals were calculated for X-linked and autosomal distributions separately. The 95% confidence intervals of the median of each M:F expression ratio distribution were calculated as described above. An ANOVA was used to test for significant differences between the means of the X-linked, M:F expression ratio distribution and the autosomal M:F expression ratio distribution within each species and F1 hybrid. Similarly, a Kruskal-Wallis test was used to test for significant differences between the medians of the X-linked, M:F expression ratio distribution and the autosomal M:F expression ratio distribution within each species and F1 hybrid.

## Results

### RNA Sequencing and Alignment

The sequencing effort yielded an average of 67.23 million reads per sample, though this number is skewed by one sample (COLZ x QUAD Female 2), which had 248.18 million reads (Supplementary Tables S1, S2). If this sample is excluded, the mean number of reads per sample is 60.53 million, ranging from 51.73 to 75.01 million. The mean mapping efficiency of parental libraries to their respective, species-specific reference genomes was 85.5% (min. 80.9%, max 88.7%, Supplementary Table S1). The mapping efficiency of parental species libraries increased when aligned to their respective AgamP4-based pseudo-genomes (mean 97.3%, min. 95.2%, max 99.1%, Supplementary Table S1).

The mapping efficiency of all libraries to their respective cDNA pseudo-genomes was reduced in comparison to either DNA genome alignment (mean 63.8%, min. 62.3%, max 65.2%, Supplementary Table S1). However, this reduction was uniform across species and may result from an incomplete AgamP4 gene annotation. F1 hybrid RNAseq reads were aligned to the respective bi-parental cDNA pseudo-genome for that cross. After filtering BAM alignments with ASE-Tigar, the average mapping efficiency for F1 hybrid libraries was 64.3% (min. 62.3%, max. 66.7%, Supplementary Table S2). Thus, the mapping efficiency of F1 hybrid and parental libraries to the cDNA pseudo-genomes was equivalent.

### Dosage compensation in An. coluzzii, An. arabiensis, and An. quadriannulatus

The analysis of dosage compensation can be biased when genes with very low expression are included. To overcome this challenge we calculated X:A and chromosome 2:3 gene expression ratios at increasing minimum RPKM thresholds from zero to 10. The number of genes included in the *An. coluzzii* - *An. arabiensis* and *An. coluzzii* - *An. quadriannulatus* comparison datasets do not differ substantially at any RPKM threshold (Supplementary Table S3). The medians of the X:A and 2:3 distributions were tested for significant differences at each RPKM threshold.

X:A median expression ratios approach 1.0 for male and female *An. coluzzii*, *An. arabiensis*, and *An. quadriannulatus* as the RPKM threshold increases (Supplementary Tables S4, S5), and at RPKM > 10, all 95% confidence intervals of the median exression ratios include 1.0 (Figure 1, Table 1). The same is true for 2:3 median expression ratios for males and females of all parental species (Figure 1, Supplementary Tables S8, S9). This result demonstrates that at RPKM > 10, 2:3 median expression ratios serve as an appropriate control with which to compare X:A expression in the effort to determine if dosage compensation is active in males.

**Figure 1.**
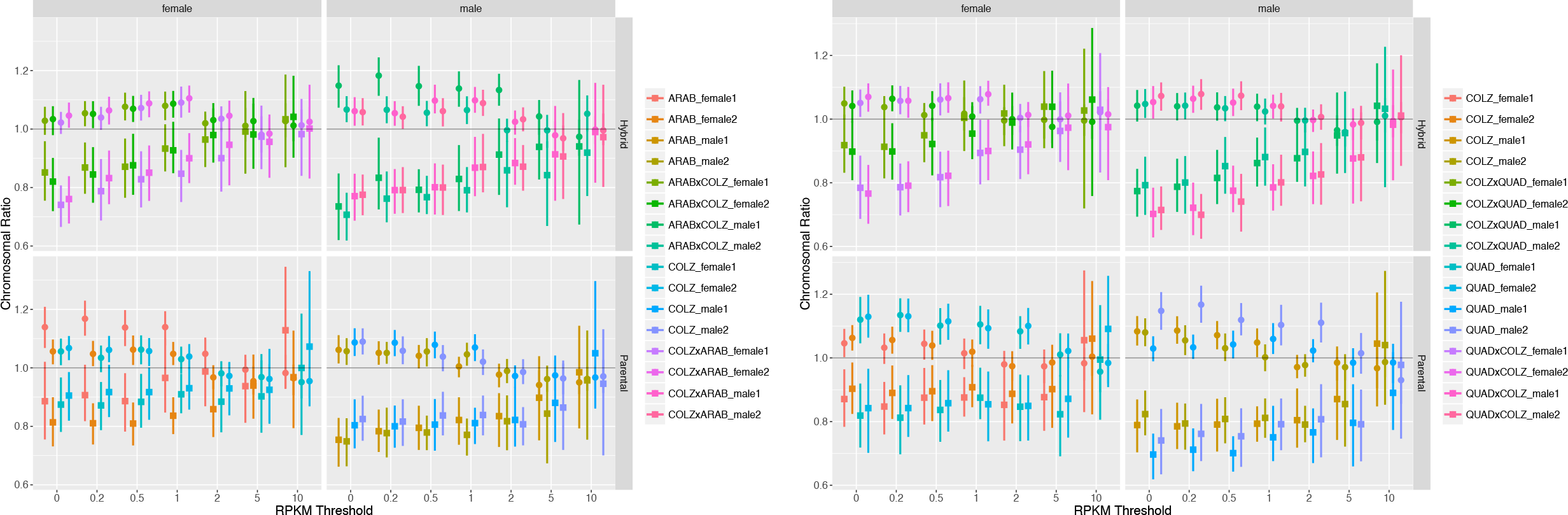
Scatter plot of X:A (squares) and 2:3 (circles) expression ratios at increasing minimum RPKM thresholds for the *An. coluzzii* - *An. arabiensis* species comparisons (left) and the *An. coluzzii* - *An. quadriannulatu*s species comparisons. Panels separate male and female (x-axis), and parental and hybrid samples (y-axis). Lines represent the 95% confidence interval of the median for each distribution.

**Table 1.** Median X:A expression ratios, and their 95% confidence intervals (CI), for each sample of the *An. coluzzii* - *An. arabiensis* species comparison and the *An. coluzzii* - *An. quadriannulatus* species comparison at RPKM > 10. The results of the between sample ANOVA (F-value and probability > F) and Kruskal-Wallis test (*X*^2^ and p-value) are reported.

| Species / F1 Hybrid | Sex | Biological Replicate | Median | 95% CI | Species / F1 Hybrid | Sex | Biological Replicate | Median | 95% CI |
| --- | --- | --- | --- | --- | --- | --- | --- | --- | --- |
| QUAD | Female | 1 | 0.96 | (0.89,1.01) | ARAB | Female | 1 | 1.13 | (0.93,1.35) |
| QUAD | Female | 2 | 0.98 | (0.92,1.04) | ARAB | Female | 2 | 0.97 | (0.79,1.13) |
| QUAD | Male | 1 | 0.99 | (0.92,1.04) | ARAB | Male | 1 | 0.99 | (0.79,1.14) |
| QUAD | Male | 2 | 0.93 | (0.86,1) | ARAB | Male | 2 | 0.96 | (0.75,1.13) |
| COLZ | Female | 1 | 0.98 | (0.93,1.03) | COLZ | Female | 1 | 1.00 | (0.77,1.19) |
| COLZ | Female | 2 | 1.00 | (0.94,1.07) | COLZ | Female | 2 | 1.07 | (0.87,1.33) |
| COLZ | Male | 1 | 0.97 | (0.9,1.03) | COLZ | Male | 1 | 1.05 | (0.86,1.3) |
| COLZ | Male | 2 | 0.99 | (0.94,1.03) | COLZ | Male | 2 | 0.95 | (0.7,1.13) |
| QUAD x COLZ | Female | 1 | 1.02 | (0.97,1.09) | ARAB x COLZ | Female | 1 | 1.03 | (0.87,1.19) |
| QUAD x COLZ | Female | 2 | 1.02 | (0.96,1.06) | ARAB x COLZ | Female | 2 | 1.04 | (0.9,1.18) |
| QUAD x COLZ | Male | 1 | 0.99 | (0.94,1.04) | ARAB x COLZ | Male | 1 | 0.94 | (0.67,1.17) |
| QUAD x COLZ | Male | 2 | 1.01 | (0.96,1.05) | ARAB x COLZ | Male | 2 | 0.92 | (0.77,1.06) |
| COLZ x QUAD | Female | 1 | 1.00 | (0.95,1.05) | COLZ x ARAB | Female | 1 | 0.98 | (0.84,1.1) |
| COLZ x QUAD | Female | 2 | 0.99 | (0.94,1.06) | COLZ x ARAB | Female | 2 | 1.00 | (0.83,1.15) |
| COLZ x QUAD | Male | 1 | 0.99 | (0.93,1.04) | COLZ x ARAB | Male | 1 | 0.99 | (0.82,1.16) |
| COLZ x QUAD | Male | 2 | 1.01 | (0.96,1.07) | COLZ x ARAB | Male | 2 | 0.97 | (0.8,1.15) |
| ANOVA F-value |  |  | 1.83 |  | ANOVA F-value |  |  | 0.38 |  |
| ANOVA Pr(>F) |  |  | 0.03 |  | ANOVA Pr(>F) |  |  | 0.984 |  |
| Kruskal-Wallis $X^2$ | | | 31.55 | | Kruskal-Wallis $X^2$ | | | 5.34 | |
| Kruskal-Wallis p-value |  |  | 0.01 |  | Kruskal-Wallis p-value |  |  | 0.989 |  |

We tested for significant differences between the medians of the X:A and 2:3 expression ratio distributions in males and females of each parental species and found no significant differences at a minimum RPKM > 10.0 (*X*^*2*^ p-value > 0.05, Supplementary Tables S8, S9). This result was confirmed through pair-wise tests for differences between of X:A and 2:3 median expression ratios between males and females, with in each parental species. X:A an 2:3 median expression ratios did not differ between any pair-wise comparisons of males and females at RPKM > 2.0 (Supplementary Tables S10, S11).

Another way to explore whether dosage compensation is operating is by comparing directly the median male to female expression ratios of X-linked and autosomal genes. Due to the under-representation of male-biased on the X-chromosome (Magnusson *et al.*, 2011, Magnusson *et al.*, 2012), we expect the male to female expression ratios of X-linked genes to be female-biased, but nonetheless close to one. Based on the findings of the chromosome ratio analyses (above), only genes with a minimum RPKM > 10.0 across all samples in a species comparison were included in this analysis.

The median M:F expression ratios ranged between 0.89 and 0.99 for X-linked genes in parental species, and between 0.91 and 1.05 for autosomal loci (Table 2, Figure 2). An ANOVA found no significant difference between the means of the X-linked and autosomal distributions, however, a Dunn’s test found significant differences between the medians of the distributions (Table 2). Despite these differences, all values are close to 1.0. In each species, the median of the M:F expression ratio distribution was lower for X-linked genes vs. autosomal genes (Figure 2), indicating, as expected, a higher proportion of male-biased genes expressed on the autosomes compared to the X chromosome. An additional explanation for this pattern (though not mutually exclusive) is a higher proportion of female-biased genes on the X chromosome relative to the autosomes. However, previous studies found that genes with female-biased expression in *An. gambiae* are randomly distributed amongst the X chromosome and autosomes (Magnusson *et al.*, 2012).

**Figure 2.**
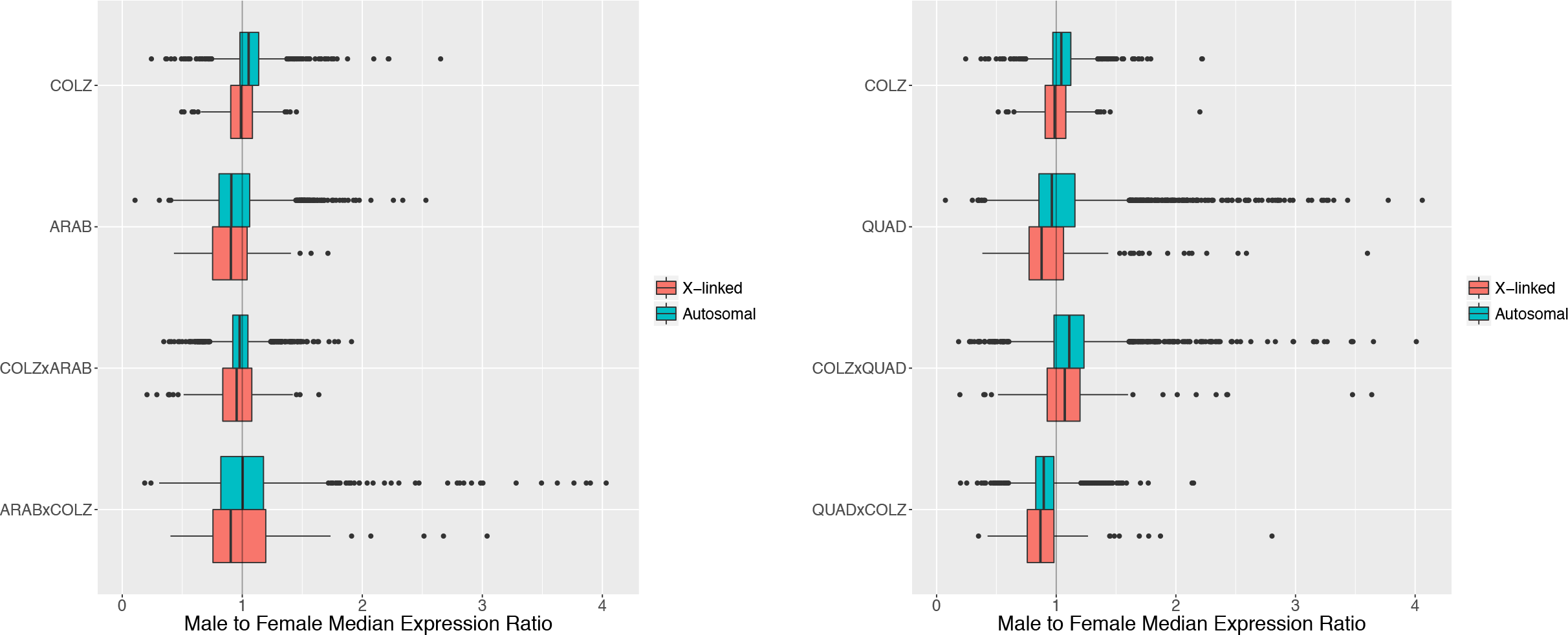
Box plot of M:F expression ratios distributions for the *An. coluzzii* - *An. arabiensis* species comparisons (left) and the the *An. coluzzii* - *An. quadriannulatus* species comparisons (right), separated by X-linked (pink) and autosomal genes (blue).

**Table 2.** Comparison of M:F expression ratio distributions, separated by X-linked and autosomal genes. Genes are only included if they have an expression level greater than 10.0 RPKM in all samples within a comparison (*An. coluzzii* - *An. arabiensis* or *An. coluzzii* - *An. quadriannulatus*). F-values and p-values (Pr(>F)) are reported for an ANOVA, which was used to test for significant differences between the means of the X-linked and autosomal M:F expression ratio distributions. *X*^2^ and p-values are reported for the Dunn’s test, which was used to test for significant differences between the medians of the X-linked and autosomal M:F expression ratio distributions.

| Species / F1 Hybrid | X-linked |  | Autosomal |  | ANOVA |  | Dunn's Test |  |
| --- | --- | --- | --- | --- | --- | --- | --- | --- |
| | Median | 95% CI | Median | 95% CI | F-value | Pr(>F) | $\chi^2$ | p-value |
| COLZ | 0.99 | (0.97,1) | 1.05 | (1.05,1.06) | 0.09 | 0.904 | 47.29 | 0.000 |
| ARAB | 0.91 | (0.87,0.95) | 0.91 | (0.9,0.92) | 0.05 | 0.997 | 6.79 | 0.005 |
| COLZ x ARAB | 0.95 | (0.93,0.97) | 0.98 | (0.97,0.98) | 0.04 | 0.999 | 7.45 | 0.003 |
| ARAB x COLZ | 0.90 | (0.84,0.95) | 1.00 | (0.99,1.02) | 0.07 | 0.968 | 6.59 | 0.005 |
| COLZ | 0.99 | (0.97,1) | 1.04 | (1.04,1.05) | 0.08 | 0.993 | 35.16 | 0.000 |
| QUAD | 0.89 | (0.83,0.94) | 0.96 | (0.95,0.97) | 0.05 | 0.999 | 25.54 | 0.000 |
| COLZ x QUAD | 1.07 | (1.03,1.11) | 1.11 | (1.1,1.12) | 0.10 | 0.976 | 10.09 | 0.001 |
| QUAD x COLZ | 0.87 | (0.85,0.89) | 0.90 | (0.89,0.9) | 0.05 | 0.999 | 13.00 | 0.000 |

These results confirm that dosage compensation is acting to balance X chromosome and autosomal gene expression in hemizygous males of *An. coluzzi*, *An. arabiensis*, and *An. quadriannulatus*, and that X chromosome dosage is equivalent between males of these species.

### Dosage compensation in Anopheles gambiae complex hybrids

Like parental species, X:A and 2:3 median expression ratios approach 1.0 for male and female F1 hybrids as the RPKM threshold increases regardless of the parental species or direction of the cross (Supplementary Tables S4, S5). At RPKM > 10 the 95% confidence intervals of the X:A median expression ratios include 1.0 for all hybrids (Figure 1, Table 1). The same is true for 2:3 median expression ratios (Supplementary Tables S6, S7).

The X:A median expression ratio of bi-directional hybrids between *An. coluzzii* and *An. arabiensis* do not differ significantly from parental species at RPKM > 0.2 (Supplementary Table 4) or at RPKM > 0.5 for 2:3 expression ratios (Supplementary Table 6). A Kruskal-Wallis test found no significant differences between median X:A and 2:3 expression ratios within both males and female hybrids at RPKM > 10 (Supplementary Table S8). Additionally, between sex comparisons of median X:A and 2:3 expression ratios in F1 hybrids found no significant differences at RPKM > 10 (Supplementary Table S10).

Similar patterns were found in bi-directional hybrids between *An. coluzzi* and *An. quadriannulatus*. The median X:A expression ratios of F1 males and females did not differ from parental species at RPKM > 10 (Supplementary Table S5), and median 2:3 expression ratios did not differ above RPKM > 0.5 (Supplementary Table S7). No significant differences were found between median X:A and 2:3 expression ratios within both males and female hybrids at RPKM > 10 (Supplementary Table S9), with on exception (COLZ x QUAD Female 1, Kruskal-Wallis p-value = 0.036). However, this difference is not significant after Bonferroni correction, and this result has no relevant bearing on our dosage compensation analyses. Finally, between sex comparisons of median X:A and 2:3 expression ratios in F1 hybrids found no significant differences above RPKM > 0.5 (Supplementary Table S11).

The median M:F expression ratios of ranged between 0.87 and 1.07 for X-linked genes in hybrids, and between 0.90 and 1.11 for autosomal loci (Table 2, Figure 2). Corresponding to the analysis of parental species, an ANOVA found no significant difference between the means of the X-linked and autosomal distributions, but a Dunn’s test found significant differences between the medians (Table 2).

These results demonstrate that hybridization between *An. coluzzii* and *An. arabiensis*, and *An. coluzzii* and *An. quadriannulatus*, does not result in the mis-regulation and disruption of dosage compensation in F1 hybrid males.

## Discussion

We analyzed dosage compensation in the closely related species within the *Anopheles gambiae* complex; *An. coluzzii*, *An. arabiensis*, and *An. quadriannulatus*, as well as in their F1 hybrids. Dosage compensation was measured by comparing the ratio of male to female gene expression on the X-chromosome and autosomes, and by comparing X-chromosome vs. autosome median gene expression levels in males and females. By analyzing bi-directional crosses between *An. coluzzii* - *An. arabiensis* and *An. coluzzii* - *An. quadriannulatu*, we tested if the species from which the Y or X chromosome is inherited affects dosage compensation in male hybrids. Specifically, these comparisons tested if the observed variation in the M locus between these closely related species may impact dosage compensation and if trans-acting factors from the heterozygous autosomes of hybrids affected X chromosome transcription differently depending on which parental species the X chromosome was inherited from.

Our results correspond with previous studies that compared X:A and M:F expression ratios in *An. stephensi* and *An. gambiae s.s.* to demonstrate that dosage compensation acts in males through the hyper-transcription of the hemizygous X (Jiang *et al.*, 2015, Rose *et al.*, 2016). Jiang *et al*. (2015) reported median X:A expression ratios in *An. stephensi* males and females between 0.92 and 0.98 at minimum RPKM thresholds between 0.0 and 4.0, with no significant differences between median X and autosomal expression in males or females above a minimum RPKM > 2.0. In addition, these authors compared expression ratios between *An. stephensi* X-linked genes and their one-to-one orthologs in *Ae. aegypti*, a species with homomorphic sex chromosomes. The median RPKM ratio of X-linked *An. stephensi* genes to their *Ae. aegypti* orthologs was close to one in both sexes after normalization, indicating full dosage compensation of these genes on the hemizygous X chromosome of *An. stephensi* males.

The evolution of sex determination and heteromorphic sex chromosomes evolved independently between *Drosophila* and *Anopheles* mosquitoes. *Drosophila* and *Anopheles* differ in their mechanisms of sex-determination (dosage vs. the *Yob* M-locus), and the mechanisms controlling dosage compensation in *Anopheles* differ from those in *Drosophila*. *Sxl* controls sex-determination in *Drosophila*, however, the *Anopheles gambiae* homolog does not have sex-specific transcripts (Saccone *et al.*, 2002), and is not involved in sex determination (Hall *et al.*, 2016). Additionally, while the orthologs of *Drosophila* MSL complex proteins are transcribed in *An. gambiae*, they are highly divergent, and their role, or lack thereof, in dosage compensation is unknown (Rose *et al.*, 2016).

The independent evolution of sex chromosomes and dosage compensation within the Culicidae (Biedler and Tu, 2016), and the rapid evolution of the M-locus in the *An. gambiae* species complex (Hall *et al.*, 2016), left open the question whether dosage compensation is disrupted in *An. gambiae* complex hybrids. Our analysis of median X:A and 2:3 gene expression ratios showed that dosage compensation is not disrupted in *An. coluzzi* - *An. arabiensis* or *An. coluzzii* - *An. quadriannulatus* hybrids, irrespective of the direction of the cross. Median X:A gene expression ratios did not differ between males and females of these hybrids; all were near 1.0, indicating complete compensation.

These results demonstrate that genes on the male X chromosome are hyper-expressed to match (on average) the expression of autosomal genes. Median male to female expression ratios for X-linked and autosomal loci differ significantly within all samples in this study (parental species and hybrids) but both of these ratios are near one. This finding is in agreement with prior work on dosage compensation in *An. gambiae s.s.* and *An. stephensi* (Rose *et al.*, 2016, Jiang *et al.*, 2015) which showed female-biased expression of X linked loci. In all species and hybrids in the present study, median M:F expression ratios are lower (female-biased) for X-linked genes compared to autosomal loci. This pattern could result from a number of evolutionary pressures that we discuss below.

Rose *et al.*’s (2016) analysis of dosage compensation in *An. gambiae s.s.* (M form) larvae and pupae found evidence for complete dosage compensation in both developmental stages. Median X:A expression ratios were found to be close to one in male and female larvae and pupae. While they note differential abundances of genes between the X chromosome and autosomes that are sex-biased in their expression, autosomal genes in this study were approximately equally expressed in both sexes, and X-linked genes were slightly female biased. Female-biased expression of X-linked genes was strongest in larvae compared to the pupae.

Female-biased expression of X-linked genes could result from an absence of dosage compensation in male testes (Rose *et al.*, 2016, Baker and Russell, 2011), an under-representation of male-biased genes on the X chromosome (Magnusson *et al.*, 2012), X chromosome inactivation in the male germline (Magnusson *et al.*, 2012), tissue-specific dosage effects in males (Baker and Russell, 2011), or a combination of these factors. In *An. gambiae s.s.* genes with testes-specific expression are under-represented on the X chromosome, while genes with ovary-specific expression are not (Baker and Russell, 2011). Furthermore, male-biased, somatically expressed genes are under-represented on the X chromosome (Baker *et al.*, 2011, Magnusson *et al.*, 2012).

Spermatogenesis occurs primarily during the 4th instar larvae and pupal stages in *Anopheles* mosquitoes (Clements 1992, Krzywinska and Krzywinski, 2009) and no genes that show male-biased expression during these developmental stages are X-linked (Magnusson *et al.*, 2011). An analysis of gene expression in the *An. gambiae* male pupae germline performed by Rose *et al* (2016) suggests that, like *Drosophila* and other Diptera (Meiklejohn and Presgraves, 2012, Vicoso and Bachtrog, 2015), the X chromosome is not hyper-expressed in the testes, and thus does not show dosage compensation in this tissue (Rose *et al.*, 2016, Baker and Russell, 2011). Sterile F1 male hybrids in the *An. gambiae* complex exhibit atrophied testes, malformed sperm, or a total lack of sperm due to the disruption of meiosis (Slotman *et al.*, 2004). However, their somatic tissues appear to develop normally.

The goal of our analysis was to confirm that dosage compensation takes place in the closely related members of the *An. gambiae* species complex, and to discern whether or not dosage compensation is mis-regulated or otherwise disrupted in F1 hybrid males. Our results show female-biased expression of X-linked loci in both hybrids and pure species males, and no significant differences in X:A and 2:3 median expression levels between hybrid and pure species males. These results indicate that dosage compensation behaves normally in hybrid males: non-sex-biased X-linked genes are hyper expressed in somatic tissues and testes have 1X expression of non-sex biased, X-linked genes.

The rapid radiation of species in the *An. gambiae* species complex provides a unique study system for investigating fundamental questions in evolutionary biology and genetics. Despite ongoing introgression and low levels of genetic differentiation between members of the *An. gambiae* complex (Fontaine *et al.*, 2015), reproductive and ecological isolation has allowed for local adaptation, behavioral divergence, and structural reorganization of their genomes (Neafsey *et al.*, 2015). This study demonstrates that despite the genetic and ecological divergence between *An. gambiae* complex member species, dosage compensation operates in pure species and hybrid males to balance X-linked and autosomal gene expression.

## Acknowledgements

This study was funded by a National Science Foundation Doctoral Dissertation Improvement Grant (award # 1601675) to K.C. Deitz and M.A. Slotman and a Texas A&M University Genomics Seed Grant to K.C. Deitz and M.A. Slotman.

